# A Novel Computational Machine Learning Pipeline to Quantify Similarities in Three-Dimensional Protein Structures

**DOI:** 10.1101/2024.08.14.607969

**Authors:** Shreyas U. Hirway, Xiao Xu, Fan Fan

## Abstract

Animal models are widely used during drug development. The selection of suitable animal model relies on various factors such as target biology, animal resource availability and legacy species. It is imperative that the selected animal species exhibit the highest resemblance to human, in terms of target biology as well as the similarity in the target protein. The current practice to address cross-species protein similarity relies on pair-wise sequence comparison using protein sequences, instead of the biologically relevant 3-dimensional (3D) structure of proteins. We developed a novel quantitative machine learning pipeline using 3D structure-based feature data from the Protein Data Bank, nominal data from UNIPROT and bioactivity data from ChEMBL, all of which were matched for human and animal data. Using the XGBoost regression model, similarity scores between targets were calculated and based on these scores, the best animal species for a target was identified. For real-world application, targets from an alternative source, i.e., AlphaFold, were tested using the model, and the animal species that had the most similar protein to the human counterparts were predicted. These targets were then grouped based on their associated phenotype such that the pipeline could predict an optimal animal species.

## Introduction

Animal models are instrumental in drug development, allowing for evaluation of target biology, pharmacology, and human safety (Morton 1998; Namdari et al. 2021). Therefore, selecting the correct animal species for analysis is crucial. Standard species selection approaches involve using legacy species based on years of previous usage, species availability or other practical considerations. When focusing on the target biology, comparing the target sequence and structure homology between human and animal species is often the first step in the process (Prior et al. 2020). Similar characteristics between the animal and human targets typically elicit similar outcomes. Therefore, target safety and efficacy assessments in animal models can determine the candidate response in a human model (Prior et al. 2020). Given the importance of animal species, further research into protein comparative analysis is vital to ensure that the optimal animal model is used in experimental studies of drug efficacy and toxicity.

Extensive research has been conducted in comparing protein structures (AlQuraishi 2021; Jain et al. 2009; Varadi et al. 2022). To address protein similarities, pairwise comparison for protein sequences is performed, yielding quantitative values such as % identities to allow for easy comparison. Modeling assessments have developed distance and contact-based measurements to directly compare protein structures (Kufareva and Abagyan 2012). The Zhang et al. group developed the Template-Modeling (TM) score model that utilizes a novel rotation matrix that focuses on local alignment of protein structures, thus resulting in faster and more accurate alignment between protein pairs in comparison to current standards (Zhang and Skolnick 2004; 2005). While such approaches demonstrate quantitative efforts, there is other valuable, biologically relevant information that is not factored into comparisons (Pearson and Sierk 2005). Additionally, even though these methods provide the tools necessary for protein comparison, they do not offer a comprehensive, end-to-end pipeline to compare a human target with multiple animal homologues to find the optimal species. Therefore, there remains a gap in the field specifically aimed at utilizing biological information in quantitative protein structure analysis that can be directly applied to species selection.

Structural data used in protein comparison is hosted in databases such as Protein Data Bank (PDB) which houses over 130,000 experimentally derived 3-Dimensional (3D) protein structures that are gathered through crystallography, and electron microscopy (Burley et al. 2017). Further efforts have been made to capture and store additional proteins efficiently by using powerful machine learning (ML) methods such as Alphafold-2 (AF2) and the AF2 database, which contains over 350,000 predicted structures and provides much more comprehensive species coverage, especially for uncommon folds (Jumper et al. 2021; Varadi et al. 2022).

Since proteins bearing high structural identities tend to have similar potencies, animal proteins are often tested for their potencies during drug development to inform protein similarity (Prior et al. 2020). Thus, bioactivity data can be used to benchmark structural similarities. For this purpose, the ChEMBL database contains 5.4 million curated bioactivity measurements for various compounds, assays, and targets (Gaulton et al. 2012). ChEMBL offers measurements such as IC50 (half-maximal inhibitory concentrations), EC (effective concentrations), and Ki (inhibitor constant) etc., (Copeland et al. 1995; Sebaugh 2011). IC50 values are valuable in assessing drug potency and response against targets and have been employed to group compound structures based on similar values (Aykul and Martinez-Hackert 2016; Sebaugh 2011; Thai and Ecker 2009).

Such structural and bioactivity data can be utilized by ML tools for predictive modeling in scientific research. When investigating proteins, ML methods have been implemented to assess protein interactions. One area of focus is Protein-Protein Interactions (PPIs), where the presence/absence of interactions between protein pairs is predicted using ensemble classifiers. The Chen et al. study utilized a gradient boosting method called Extreme Gradient Boosting (XGBoost) for dimensionality reduction and biological feature selection (Chen et al. 2020). XGBoost is a tree-based model for solving classification and regression-based problems. Its advantages include the ability to scale, handle large or sparse datasets, and accommodate instance weights in approximate tree learning to create an end-to-end system (Chen and Guestrin 2016). The XGBoost-based ensemble classifier in Chen et al. yielded a higher PPI prediction accuracy for multiple biological datasets in comparison to classifiers such as linear regression and support vector machines (SVMs) (Chen et al. 2020). Furthermore, to better understand biological activity, classifiers such as random forests and SVMs have been applied to transform gene-based cluster data and predict antibacterial/antifungal activity in natural products (Walker and Clardy 2021).

In the context of species selection, valuable information from biologically relevant tertiary structures is not utilized to inform protein similarities. In this study, we develop an end-to-end computational pipeline to identify animal proteins that are most similar to their human counterparts. This quantitative approach can objectively compare protein homologues. We compile collections of protein targets gathered from bioactivity and structural databases, which then generate protein structure and function-based attributes. With IC50 values used to benchmark protein similarity, regression analysis is performed to predict the optimal animal homologue for human proteins. Model predictions for proteins are grouped by their associated phenotype to identify the optimal animal species for investigating that phenotype.

## Methods

The computational pipeline to quantify homology for protein tertiary structures is detailed in Figure 1. We began with the dataset curation and generation module (Figure 1.A), which centered on processing protein entries acquired from ChEMBL. Target protein entries were extracted and processed based on multiple selection criteria (Figure 1.A.1). The Uniprot IDs of the final set of filtered entries were converted to PDB IDs. Then, a new dataset of entries, detailed in Figure 1.A.2, was generated which paired human target entries with the animal homologues available for those targets. Each entry in this structural dataset also incorporated IC50 activity values, which underwent a normalization process, resulting in similarity scores that were used to compare the human targets to their animal counterparts. The newly generated data was analyzed in the Machine Learning Module (Figure 1.B). Structural alignment packages, and natural language processing techniques such as topic modeling were used to generate features for each entry in the structural dataset (Figure 1.B.1). For the machine learning model, two datasets of features were generated, one based on data from the Research Collaboratory for Structural Bioinformatics (RCSB) database (Rose et al. 2016), and one based on data from AF (Figure 1.B.2). A large parameter study was conducted to determine the optimal model based on the similarity scores.

**Figure 1:**
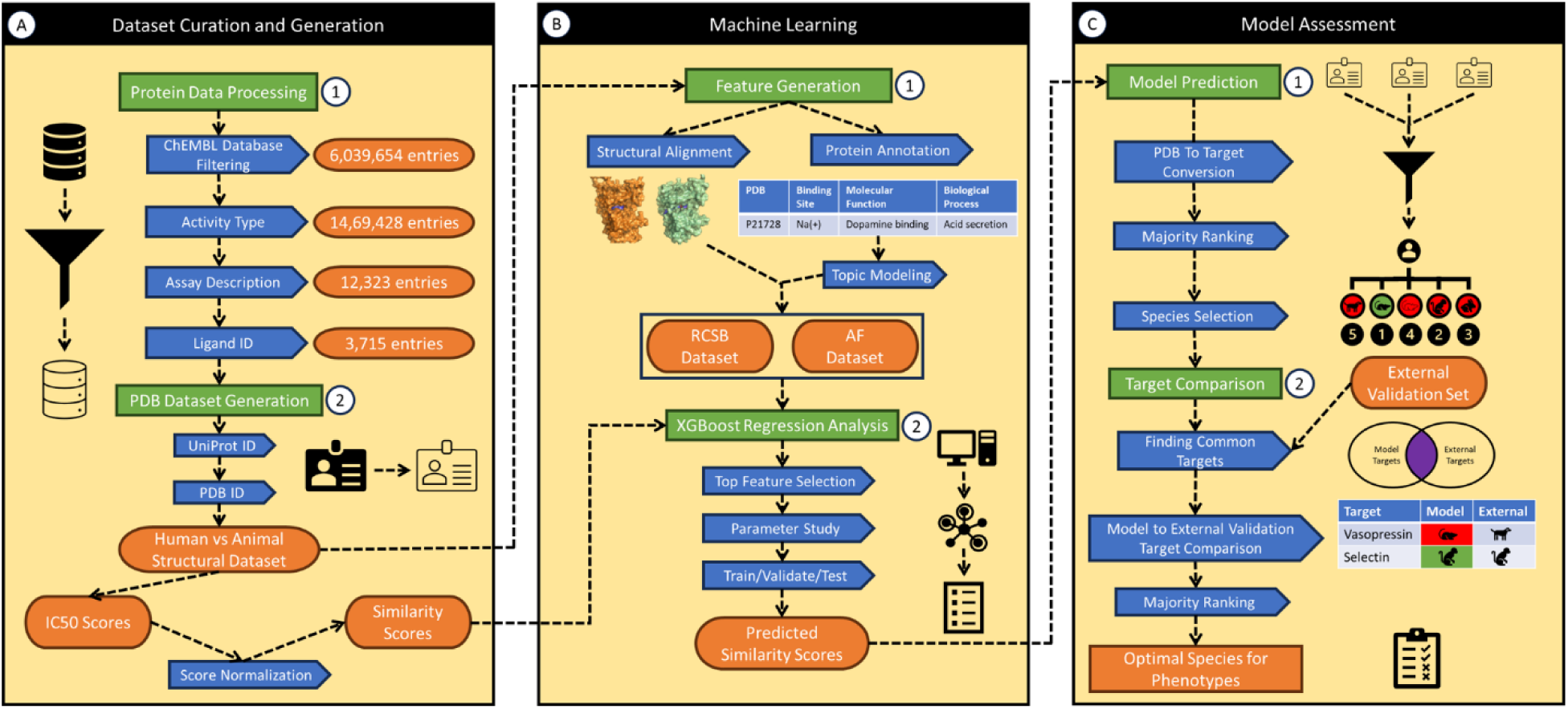
Flowchart of computational pipeline. Flowchart shows the three main modules: Dataset Curation and Generation (A), Machine Learning (B) and Model Assessment (C). Protein Data Processing (A.1) and PDB Dataset Generation (A.2) help generate the human vs animal structural dataset. This dataset is processed through Feature Generation (B.1) and used in XGBoost Regression Analysis (B.2). The resultant predicted similarity scores are used for Model Validation (C.1) and Target Comparison (C.2).

These scores were used as ground truth by the regression model in the Model Assessment module (Figure 1.C). During model prediction, multiple PDB entries for a target protein were aggregated together (Figure 1.C.1). The various species of PDB entries were ranked, and an optimal species was selected per target during the Species Selection component. The next element of the pipeline involved comparing the model predictions to an external dataset, a real-world case study which utilized data from Off-X and Pharmapendium databases (Figure 1.C.2). Targets predicted by the model were grouped together based on their associated phenotype and compared to the real-world case study.

### Dataset Curation and Generation

ChEMBL was downloaded, which contained attributes such as protein names and IDs, associated ligands, assay descriptions, various measured bioactivities, and species of origin of proteins. With the objective to compile a set of entries for human and animal target comparison, the dataset was curated in the following way, as seen in the Protein Data Processing sub-module (Figure 1.A.1). A target’s activity type was the first filter step where entries containing IC50 data were selected. These IC50 values would be later used in the regression model. Entries were also filtered based on their assay description, where entries with the keyword “binding” were selected to incorporate binding-based assays performed on the targets. Target proteins which contained entries with the human species as well as another non-human species commonly used in animal studies such as rat, mouse, dog, rabbit, marmoset, rhesus monkey, and cynomolgus monkey, were selected such that comparisons between the human and animal entries could be performed. Since each human target protein contained data associated with multiple different ligands, the ligand name associated with each non-human entry was also matched. This ensured additional specificity during target comparisons. Once this dataset was curated, the Uniprot IDs specific to each target, species and ligand were identified. In the PDB Dataset Generation sub-module (Figure 1.A.2), these IDs were parsed in the RCSB database to find PDB entries which contained 3D protein structure data as well as other additional properties. Similarly, the Uniprot IDs were also parsed directly in the AF database to generate a set of protein structure entries. Then, entries for human-animal comparison were created. For each unique target protein and ligand combination, the human PDB entries were matched up against their non-human homolog PDB entries. Thus, human to animal structural datasets were created for both the RCSB and AF databases.

### Machine Learning

These two datasets were processed to generate features for the machine learning model (Figure 1.B). Both structural as well as annotation data were utilized to generate features (Figure 1.B.1). The TM-align package was used to facilitate a structural comparison between two PDB entries, resulting in a total of 14 metrics (Zhang and Skolnick 2005). Firstly, chain scores were calculated based on cross comparisons between human and animal proteins. A 3-coordinate translation matrix was then generated to align the centroids of the two proteins. Subsequently, a 9-coordinate rotation matrix was created to maximize the overlap between the two proteins. Additionally, a sequence alignment identity score was also calculated using the Biopython package (Cock et al. 2009). Table 1 details the description for each variable.

**Table 1:** Structural features for protein comparison. 15 total features were generated for each human to animal protein comparison. Some of metrics output multiple values as mentioned in the description. The first 14 metrics were generated by TM-align while the Identity feature was calculated using Biopython. The range of values for each variable is also provided.

| Structural Metric Name | Description | Range |
| --- | --- | --- |
| Chain Score | TM-score between human protein structure to animal protein structure relative to the human structure | [0 , 1] |
|  | TM-score between animal protein structure to human protein structure relative to the animal structure |  |
| Translation Matrix | 3-coordinate matrix for aligning the centroid of human protein structure with centroid of the animal protein structure | [-40 , 60] |
| Rotation Matrix | 9-coordinate matrix for rotating the human protein structure to maximize overlap with animal protein structure | [-1 , 1] |
| Identity | Pairwise sequence alignment between human protein structure and animal protein structure | [0 , 100] |

Furthermore, annotation data was generated for each PDB entry. In this step, each PDB entry was linked with the corresponding UniProt ID. All IDs were subsequently queried in UniprotKB and custom annotations were added from the default 14 categories. Details of the custom annotations are listed in the supplement table S1. Each of the custom annotations was selected with manual inspection to ensure that it was relevant to the current study and there existed sufficient information for the queried ID list. Given that the annotations contained text data, a natural language processing approach was taken to derive features from each annotation entry. Using topic modeling, text from each of the 14 categories were inputted into a Latent Dirichlet Allocation (LDA) model which grouped the entries based on tokens or words (Blei et al. 2003). Each category had a model which grouped the PDB entries into 3 distinct categories based on the topic words generated in each of the category. Thus, each individual human and animal PDB entry had a rank of 1, 2 or 3 for each category. Given that some categories were missing data for certain PDB entries, a total of six categories were selected for regression analysis: protein families (PF), function, sequence similarities (SS) and gene ontology (GO) entries for biological process (GOBP), cellular component (GOCC) and molecular function (GOMF). After generating the ranking data, the One-hot-encoding (OHE) technique was used to convert each categorical feature into a numerical feature, where binary notations were used to denote the presence or absence of the PDB entry in the corresponding category, resulting in three features per category (Draper and Smith 1998; Kaplan 2021). As a result, the annotation data from the 6 categories were broken down into OHE entries. This created a total of 18 features per PDB entry, resulting in a total of 36 features for each human to animal PDB comparison. Together, the structural and annotation data combined for a total of 51 features for each human to animal PDB comparison.

To set up the following machine learning procedure, truth labels were created for each comparison using IC50 values. IC50 activity scores were extracted for each PDB by locating the Uniprot ID associated with the PDB. For each Uniprot ID, there are multiple entries in the RCSB database due to various ligands and assays associated with each entry. To standardize multiple values for one Uniprot ID, the IC50 values between the 25% and 75% of the entire range were used to calculate a standardized average IC50 value for each PDB entry. For human to animal comparisons, a normalized IC50 value was calculated, as shown by (1-2), where *X*_*i*_ is the initial IC50 value, *L*_*i*_ is the log-transformed value, *L*_*H*_ is the log-transformed value of the human species and *X*_*N*_is the normalized IC50 value. Once the normalized values were calculated, the minimum value was subtracted from the entire set to create a new set of values denoted *X*_*S*_, as seen in (3) and then divided by the maximum, as shown by (4) to calculate a final version of “Similarity Scores” denoted *X*_*SS*_ with the range of 0 to 100. Due to this data transformation, scores for entries closest to human were shifted rightward, where a score closest to 68.40905 was determined as the closest value to a human entry. Thus, when looking at entries for multiple species, the species with the similarity score closest to the human value would be deemed as the optimal species.

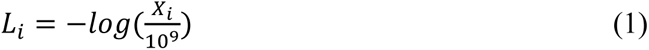

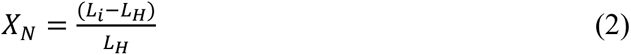

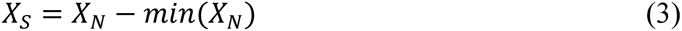

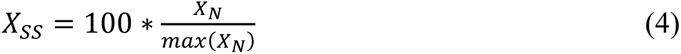

In this pipeline, two separate datasets were created. The RCSB dataset was reserved for training/validation/testing purposes to develop the regression model while the AF dataset was used to make final predictions on the protein targets. For regression analysis, models such as CatBoost and XGBoost were initially tested and XGBoost was deemed as the optimal choice based on its performance. The XGBoost regression model was developed using the annotation and alignment features to predict similarity scores for a human and animal PDB entry comparison (Figure 1.B.2). This model was setup in three parts. The model was initially run to identify top contributing features in the dataset. Then, a similarity score was predicted for each entry by the model.

Next, a large parameter optimization step was conducted to determine the best parameter set that increased the regression model’s performance. Metrics were calculated for the training, validation, and testing RCSB set. A Performance Metric (*P*_*M*_) designed to assess the predicted similarity score’s closeness to the truth score as well as the human score was generated. Equation (5) shows the equation which was derived from the Mean Absolute Error (MAE) formula (Wang and Lu 2018), where *S*_*p*_ denotes the predicted score, *S*_*t*_ denotes the truth score and *H*_*t*_ denotes the human truth value. Given that the model’s predicted similarity scores being closer to the true scores as well as the human value is deemed to be optimal, a *P*_*M*_ value closer to 0 signifies a higher model performance. Each parameter set was run three times, and the recorded metrics were averaged. Some notable variables varied in this optimization process include XGBoost specific parameters such as alpha, lambda, step size, tree depth, subsample ratio, column subsampling value, number of important features, number of parallel trees, and size of the testing set.

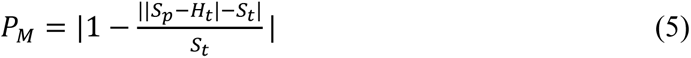

The XGBoost model developed using the best parameter set based on the RCSB testing set’s *P*_*M*_ value was used to generate similarity score predictions for entries in the AF set.

### Model Assessment

Model performance was assessed using the similarity score predictions (Figure 1.C). Animal homolog entries for each unique human entry were identified in the AF dataset (Figure 1.C.1). Next, the species with the closest similarity score to the human score was determined and majority ranking was performed on all the entries for a target such that the animal species with the highest number of entries would be deemed as the optimal species for a target.

A phenotype-based dataset was gathered to serve as a real-world case study. This dataset was compiled using integrated data from Pharmapendium and Off-X databases, where reported adverse events, referred to as phenotypes, of preclinical and clinical studies were documented. From these databases, drugs for which the phenotype of interest was observed in both preclinical and clinical species were defined as true positives. The pharmacology for these drugs, i.e., intended, and unintended target proteins, was curated from DrugBank (Wishart et al. 2018). The resulting targets were compared with the AF dataset and the overlapping target proteins were tested using our model. Further filtering within the targets was performed by ensuring that the animal species available in the case study existed within the regression model’s total list of species. Then, majority ranking was performed on all the targets in one phenotype to determine the optimal species for the phenotype based on the model predictions. These results were then compared to the real-world dataset to determine if for a given phenotype, the model’s predicted species agreed with the species deemed as most translatable or to be exhibiting the closest resemblance to the human model based on literature curation.

## Results

### Preparation of Databases

Throughout this study, we developed multiple datasets for protein structure comparison. Firstly, the ChEMBL database was curated and filtered to develop a list of human and animal entries with matching targets and ligands. This database of 6,039,654 entries was reduced to 1,469,428 entries when filtered based on activity type (IC50), then to 12,323 entries when screening for binding activity in the assay description and finally to 3,715 after matching ligands for each target and ensuring that each target has an entry for human species and one for a non-human species of interest. The Uniprot IDs from this curated dataset were used to find PDB file IDs which contained structural information on these targets. Based on the number of PDB files available per Uniprot ID, the number of human to animal comparison entries generated for a certain protein-ligand combination ranged from one to several hundred.

Comparison entries for two structural datasets were created, the RCSB based dataset for building the regression model, and the AF based dataset for making final predictions to compare to the real-world case study dataset. The RCSB dataset consisted of 25,089 entries. The AF dataset also included entries that intersected between the ChEMBL database and the case study dataset, where PDB entries with measurements from other activity types were also included, to expand the size of the AF dataset to 2,181 entries. We performed exploratory data analysis to visualize the distribution of entries in both these datasets. Figure 2 shows the similarity score distribution for the RCSB dataset where the blue line denotes the distribution density, and the red dashed line is the human score of 68.41 which is a benchmark used for comparing and selecting animal species. Furthermore, the distribution of the eight animal species of interest for all the human-animal comparisons are also provided as percentages in Figure 2.B and C. While the RCSB dataset primarily contained four species including Sus scrofa, Rattus norvegicus, Mus musculus and Oryctolagus cuniculus, the AF dataset contained entries for all eight species. Given that the AF dataset contained multiple activity types, a similarity score distribution based on IC50 values was not generated.

**Figure 2:**
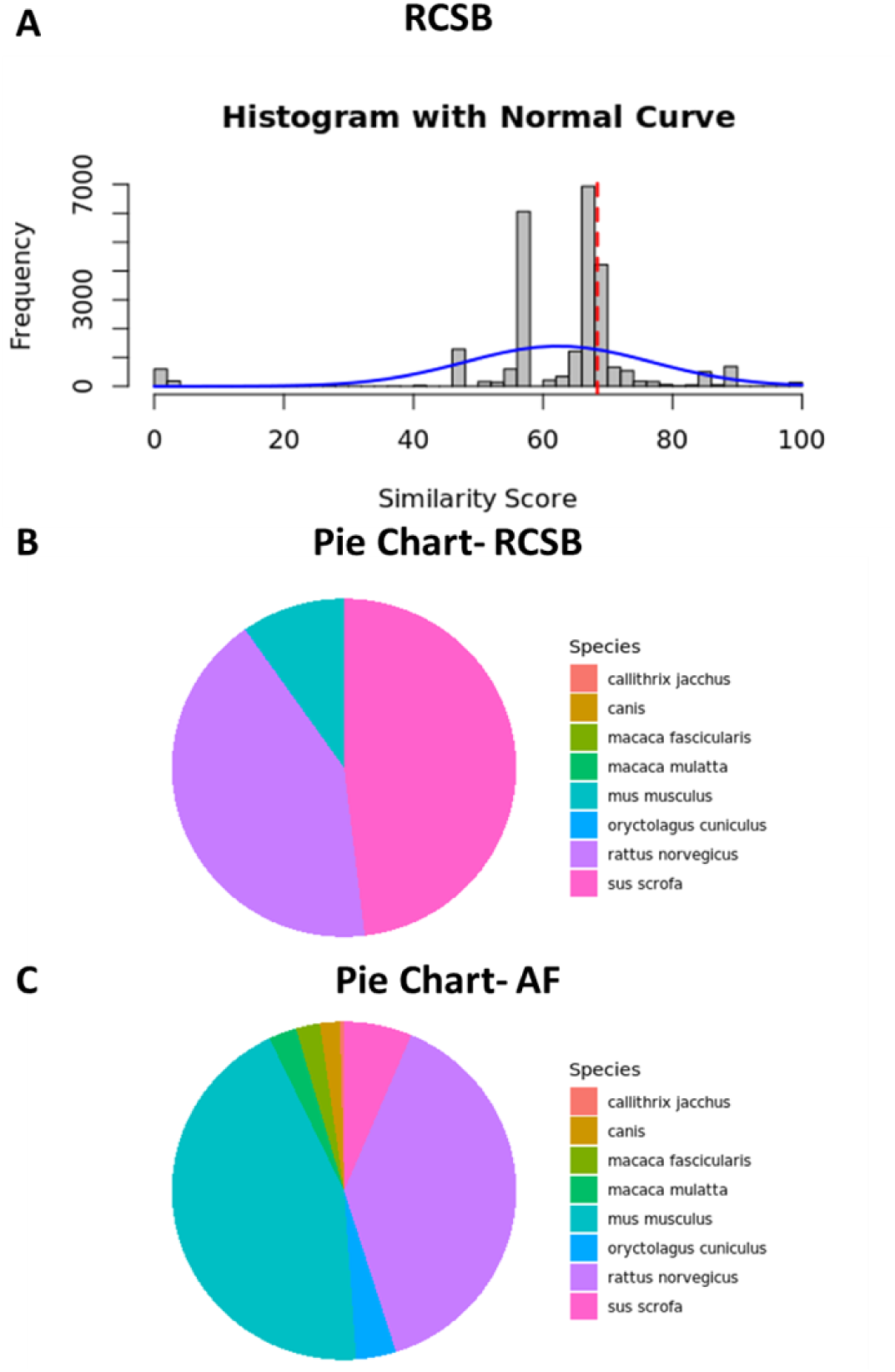
Similarity score histograms and score distribution pie charts for model datasets. (A) shows the similarity score histogram for the RCSB dataset where the blue line denotes the distribution density, and the red dashed line denotes the human score (68.41). Animal species distribution pie charts for RCSB (B) and AF (C) represent the percent distribution of the various animal species, where Rattus norvegicus, Sus scrofa and Mus musculus are amongst the most prevalent species in the datasets.

The RCSB dataset was used to develop the regression model. This set was split into 90% training and 10% testing set, where the training set was again split (90/10) to create an additional validation set. To find the best model using the *P*_*M*_ value in the testing set, a parameter study was conducted that varied several components of the XGBoost model. Figure 3 briefly highlights the summary of the parameter study where the line graph shows the effect of the number of top features, ranging from 5 to 50 in increments of 5, used in building the model on *P*_*M*_, averaged over three runs. A model developed using the top 10 features demonstrated the best *P*_*M*_ value as represented by the black dot in Figure 3. Furthermore, Table 2 shows the top 10 ranked features used by the best model, where five features are from the structural data and the other five are from the topic modeling data. Metric names such as “Topic 2” and “R5” denotes the second OHE topic metric, or the 5^th^ rotation matrix coordinate in the feature data respectively. The gain value, describing the relative contribution of each feature in the model is also provided (Chen and Guestrin 2016). Interestingly, GOMF and SS attributes contained multiple features in this set.

**Figure 3:**
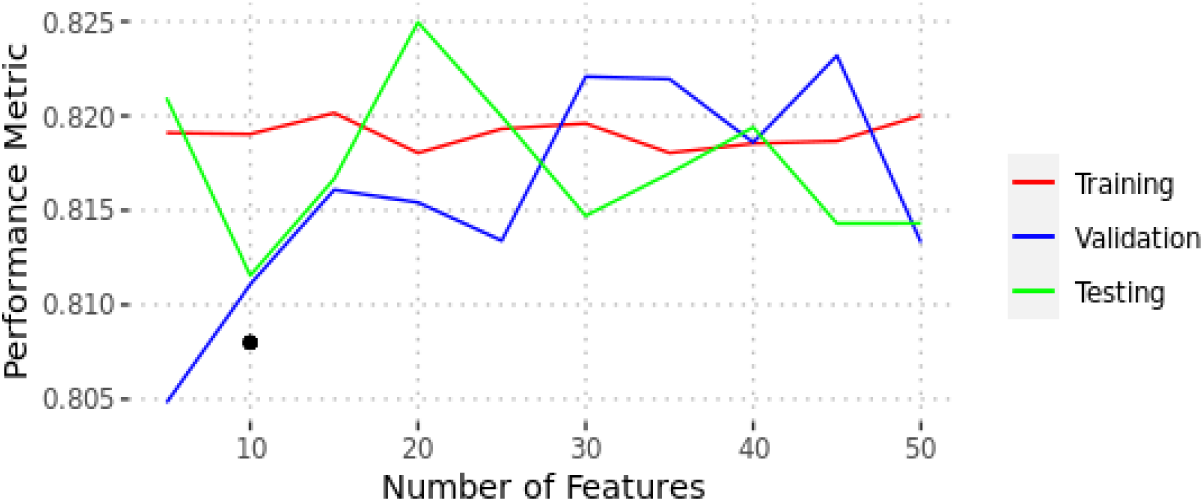
*P*_*M*_ measured across varying number of features in the model. The *P*_*M*_, averaged over three runs, when varying the number of model features from 5 to 50 is shown for the RCSB Training (Red), Validation (Blue), and Testing (Green) dataset. The black dot denotes the model with the best *P*_*M*_ value (0.808) which was developed using the top 10 features.

**Table 2:** Most important features derived from XGBoost model. The rank, description, and gain (relative contribution) of the top 10 most important features determined by the XGBoost model. Metric names such as “Topic 1” and “T2” denotes the first OHE topic metric, or the 2nd coordinate in the translation matrix in the feature data respectively.

| Rank | Metric Name | Description | Gain |
| --- | --- | --- | --- |
| 1 | Identity | Pairwise sequence alignment between human sequence and animal sequence | 0.192 |
| 2 | Animal Entry GOCC Topic 2 | GOCC annotation-based topic 2 for animal entry | 0.102 |
| 3 | Animal Entry GOMF Topic 2 | GOMF annotation-based topic 2 for animal entry | 0.0743 |
| 4 | Chain 1 | TM-score from human protein structure to animal protein structure | 0.0733 |
| 5 | Human Entry GOMF Topic 2 | GOMF annotation-based topic 2 for human entry | 0.0600 |
| 6 | Translation Matrix- T2 | 2<sup>nd</sup> coordinate in translation matrix for aligning the human protein structure onto the animal protein structure | 0.0534 |
| 7 | Rotation Matrix- R5 | 5<sup>th</sup> coordinate in rotation matrix for maximizing overlap between the human protein structure and the animal protein structure | 0.0484 |
| 8 | Human Entry SS Topic 2 | SS annotation-based topic 2 for human entry | 0.0448 |
| 9 | Chain 2 | TM-score from animal protein structure to human protein structure | 0.0415 |
| 10 | Human Entry SS Topic 1 | SS annotation-based topic 1 for human entry | 0.0347 |

### Comparison to Real-World Case Study

The model with the best parameters identified from the parameter study was used to make predictions on structures obtained from the AF dataset. A total of 419 target proteins which existed in both the model predictions and the real-world case study were selected. Additional filtering was performed to ensure that the species available in the case study was within the range of all the available species for each target in the model, resulting in a final set of 141 target proteins. Table S2 in the Supplement shows the resulting predictions from the best model, and the phenotype associated with those targets based on the real-world dataset.

The 141 targets were associated with several phenotypes. Tables 3, 4 and 5 show the model predictions for targets associated with the phenotype as well as the species that is most translatable to human based on literature curation for Hypotension, Dry Eye, and Anemia respectively (Barabino et al. 2005; Cayla et al. 2007; Gonzalez-Casas et al. 2009; Hanna et al. 2007; Kim et al. 2014; Mantelli et al. 2013; Moon et al. 2020; Oh et al. 2015; Shinomiya et al. 2018; Tanimoto et al. 1994). Our model predicted mouse as the optimal species for a majority of the targets for all three phenotypes. By using the previously mentioned majority ranking method, the optimal species for each phenotype was then established, where mouse models were predicted to be optimal for all three phenotypes. Additionally, several experimental studies in literature have utilized mouse models to investigate various targets associated with these three phenotypes (Barabino et al. 2005; Cayla et al. 2007; Gonzalez-Casas et al. 2009; Hanna et al. 2007; Kim et al. 2014; Mantelli et al. 2013; Moon et al. 2020; Oh et al. 2015; Shinomiya et al. 2018; Tanimoto et al. 1994).

**Table 3:** Model Predictions for targets associated with hypotension. For the hypotension phenotype, the most translatable species in the real-world dataset based on literature curation (Cayla et al. 2007; Tanimoto et al. 1994), as well as the model predictions for targets associated with hypotension are listed below. For 11 out of 12 targets, our model predicts mouse as the optimal species, in agreement with hypotension focused animal models found in literature.

| Phenotype | Most Translatable Species | Associated Target | Symbol | Model Prediction |
| --- | --- | --- | --- | --- |
| Hypotension | mus musculus | Renin | REN | mus musculus |
|  |  | Catechol O-methyltransferase | COMT | mus musculus |
|  |  | Solute carrier family 15 member 2 | SLC15A2 | mus musculus |
|  |  | Equilibrative nucleoside transporter 1 | SLC29A1 | mus musculus |
|  |  | Adenosine deaminase | ADA | mus musculus |
|  |  | Adenosine A1 receptor | ADORA1 | mus musculus |
|  |  | Adenosine A3 receptor | ADORA3 | mus musculus |
|  |  | Adenosine A2b receptor | ADORA2B | mus musculus |
|  |  | Adenosine A2a receptor | ADORA2A | mus musculus |
|  |  | Purine nucleoside phosphorylase | PNP | mus musculus |
|  |  | Trace amine-associated receptor 1 | TAAR1 | rattus norvegicus |
|  |  | Interleukin-2 | IL2 | mus musculus |

**Table 4:** Model predictions for targets associated with dry eye. For the dry eye phenotype, the most translatable species in the real-world dataset based on literature search (Barabino et al. 2005; Mantelli et al. 2013; Moon et al. 2020; Oh et al. 2015; Shinomiya et al. 2018), as well as the model predictions for targets associated with dry eye are listed below. Our model predicts mouse as the optimal target for all 8 targets associated with dry eye, which agrees with the experimental studies found in literature.

| Phenotype | Most Translatable Species | Associated Target | Symbol | Model Prediction |
| --- | --- | --- | --- | --- |
| Dry eye | mus musculus | Vascular endothelial growth factor receptor 2 | KDR | mus musculus |
|  |  | Vascular endothelial growth factor receptor 3 | FLT4 | mus musculus |
|  |  | Vascular endothelial growth factor receptor 1 | FLT1 | mus musculus |
|  |  | Hepatocyte growth factor receptor | MET | mus musculus |
|  |  | Tyrosine-protein kinase LCK | LCK | mus musculus |
|  |  | Tyrosine-protein kinase SRC | SRC | mus musculus |
|  |  | Tyrosine-protein kinase Lyn | LYN | mus musculus |
|  |  | Serine/threonine-protein kinase Aurora-A | AURKA | mus musculus |

**Table 5:** Model predictions for targets associated with anemia. The most translatable species in the real-world dataset based on literature search (Gonzalez-Casas et al. 2009; Hanna et al. 2007; Kim et al. 2014), as well as the model predictions for targets associated with anemia are listed below. Mouse is predicted as the optimal species for 5 out of the 6 associated with anemia. Therefore, our majority ranking-based approach is in agreement with animal model studies found in literature that focus on anemia.

| Phenotype | Most Translatable Species | Associated Target | Symbol | Model Prediction |
| --- | --- | --- | --- | --- |
| Anemia | mus musculus | Tankyrase-1 | TNKS | mus musculus |
|  |  | Thymidine kinase, cytosolic | TK1 | mus musculus |
|  |  | Tankyrase-2 | TNKS2 | mus musculus |
|  |  | Bradykinin B1 receptor | BDKRB1 | oryctolagus cuniculus |
|  |  | Ras-related C3 botulinum toxin substrate 1 | RAC1 | mus musculus |
|  |  | Leukotriene A4 hydrolase | LTA4H | mus musculus |

Furthermore, to compare our XGBoost model with current practices, sequence and structure-based models were generated to predict the optimal species for the three highlighted phenotypes. These models solely used the Identity and Chain 1 features respectively and did not utilize the XGBoost regression model. In comparison to our best model, the sequence and structure-based models predicted mouse as the optimal species for less targets in each phenotype type. Specifically for hypotension, our model predicted mouse as the optimal species in 11 out of the 12 targets. In contrast, both the sequence and structure-based models predicted mouse for only 7 targets. As such, our model provided more confidence than the sequence and structure-based models when selecting mouse models to be the optimal species for the investigating hypotension.

## Discussion

In this study, we successfully developed an ML pipeline that used structural and annotation-based information to predict overall protein similarities to inform primary species choice, benchmarked against IC50 data. Our model predicted targets from the AF-2 dataset as well as the phenotypes associated with those targets in the real-world case study.

This model takes a novel approach regarding the data used for performing protein comparison. While current methods utilize sequence-based and structural data (Kufareva and Abagyan 2012; Pearson and Sierk 2005), our pipeline also incorporates nominal data extracted by performing natural language processing on proteins’ biological information such as protein families, gene ontology etc. Furthermore, the machine learning component of this pipeline utilizes similar framework as in literature. XGBoost models have been used to define important feature data in biological studies (Zhong et al. 2018). Our model utilized XGBoost in a more end-to-end manner, where it was used to establish the top-ranking features from the structural and nominal data, and also to perform regression analysis on the dataset, using IC50 values during protein comparison.

IC50 measurements’ have been utilized to categorize compound structures with similar target response and therefore can be used to benchmark target homology comparison (Aykul and Martinez-Hackert 2016; Sebaugh 2011; Thai and Ecker 2009). Our pipeline uses IC50 values to generate similarity scores which are used as ground truths in the regression model. This successfully allows the pipeline to determine which animal homologs are closest to the human targets.

While our pipeline primarily identifies species for individual protein targets, this model can also be used to suggest optimal species to study a given phenotype based on the predictions of multiple targets contributing to the phenotype of interest. Through majority ranking, the most frequently selected animal species was chosen as the best fit to utilize for a given phenotype. Notably, for phenotypes of dry eye, hypotension and anemia, the pipeline predicted mouse as the optimal species for the majority of targets. These findings are also in agreement with literature studies that have utilized mouse models to understand several targets in these phenotypes (Barabino et al. 2005; Cayla et al. 2007; Gonzalez-Casas et al. 2009; Hanna et al. 2007; Kim et al. 2014; Mantelli et al. 2013; Moon et al. 2020; Oh et al. 2015; Shinomiya et al. 2018; Tanimoto et al. 1994). Specifically, mouse models have been used to investigate hypotension-related targets like renin, as well as anemia markers such as Tankyrase-2, which has been linked to various forms of cancer (Lehtiö et al. 2013; Tanimoto et al. 1994). Our pipeline predicts mouse models as the optimal species for both these markers, as highlighted in Table 3 and 5. Furthermore, therapeutics development for multifactorial diseases can target a multitude of proteins. In the case of dry eye, therapeutics often target growth factors or protein kinases. Inhibitor drugs often target proteins from the tyrosine kinase family, such as SRC and LCK (Coco et al. 2023). Our pipeline predicts mouse as the optimal species for both these proteins (Table 4), in good agreement with using mouse models to study dry eye, especially if the therapeutic focus is on tyrosine kinases. As such, our pipeline’s ability to predict the optimal species for individual targets can be extrapolated onto selecting the appropriate species for a phenotype by majority ranking the prediction results made from multiple contributing targets.

Given that the core of the model focuses on determining the similarity between two targets, it can have many applications in protein analysis. One such application would be off-target protein analysis. Using a known protein, other similar proteins could be identified based on their protein structures as potential off-target proteins. This might provide value for hazard identification and potentially help to form hypothesis to address a given *in vivo* phenotype.

### Limitations and Future Considerations

There are limitations to this model. Firstly, while the ChEMBL database contains a large magnitude of entries, filtering it down to match the purposes of our analysis resulted in a smaller set of entries. Each direct comparison of a specific target and ligand had a small amount of animal samples, and a select number of animal species available, which restricted the total number of targets for comparison with the real-world case study. Furthermore, not every species is tested for all the targets in the case study, limiting the range of species available for comparison. Lastly, based on the species distribution, there is a lack of cynomolgus monkey data available and an abundance of mouse and rat data, thus restricting the model’s exposure to certain species and potentially leading to a bias.

There are several considerations to make which could improve the model coverage. Specific animal PDB entries could be manually added for each target such that more species are available, and each target has more entries for comparison. Furthermore, additional structural alignment methods could be used to generate more structural metrics for the structural alignment. Finally, improving the regression model to better utilize categorical data would offer a better opportunity to incorporate the nominal data into the model.

## Supplemental Tables

**Table S1:** All selected protein entry annotations and their descriptions by category from UNIPROT database. Categories from this table were used for generating topic modeling-based features.

| Category | Entries | Details |
| --- | --- | --- |
| **Names & Taxonomy** | Entry Name | UNIPROT ID |
|  | Gene Name | All Name(s) of the gene(s) encoding the protein |
|  | Organism | Organism name |
|  | Protein Names | Name(s) and synonym(s) of the protein |
|  | Gene Names (primary) | Primary Gene Name |
| **Sequences** | Length | Length of the canonical sequence |
|  | Mass | Molecular Mass in Daltons |
| **Function** | Activity Regulation | Description of the regulation of the catalytic activity of an enzyme, transporter, transcription factor |
|  | Binding Site | Binding site for any chemical group (co-enzyme, prosthetic group, etc.) |
|  | DNA Binding | Description of the position and type of each DNA-binding domain present within the protein |
|  | Function (CC) | General description of a protein's function(s) |
|  | Site | Interesting single amino acid site on the sequence |
| **Miscellaneous** | Keywords | List of UniProtKB keywords (controlled vocabulary) |
| **Interaction** | Interact With | Cross-references to UniProtKB entries describing protein interactors, as reported in the Intact database |
| **GO** | Gene Ontology (BP) | Gene Ontology Biological Process |
|  | Gene Ontology (CC) | Gene Ontology Cellular Component |
|  | Gene Ontology (MF) | Gene Ontology Molecular Function |
| **PTM / Processing** | Post-translational Modification | Sequence position-independent description of post-translational modifications |
| **Structure** | Beta strand | Positions of the experimentally determined beta strand region(s) |
|  | Helix | Positions of the experimentally determined helical region(s) |
|  | Turn | Positions of the experimentally determined Turn region(s) |
| **Family & Domains** | Coiled coil | Positions of region of coiled coil |
|  | Motif | Short (up to 20 amino acids) sequence motif of biological interest |
|  | Protein families | Name of family(ies) to which the protein belongs to |
|  | Sequence similarities | Sequence position-independent annotation (comments) |
|  | Zinc finger | Position(s) and type(s) of zinc fingers within the protein |
|  | Domain (CC) | Type (and number) of domains in the canonical sequence, as listed in the 'General annotation (Comments)' section |
|  | Domain (FT) | Position and type of each modular protein domain |
| **Chemistry** | DrugBank | DrugBank IDs and name |
| **Enzyme & Pathway** | Reactome | Reactome IDs and name |

**Table S2:**
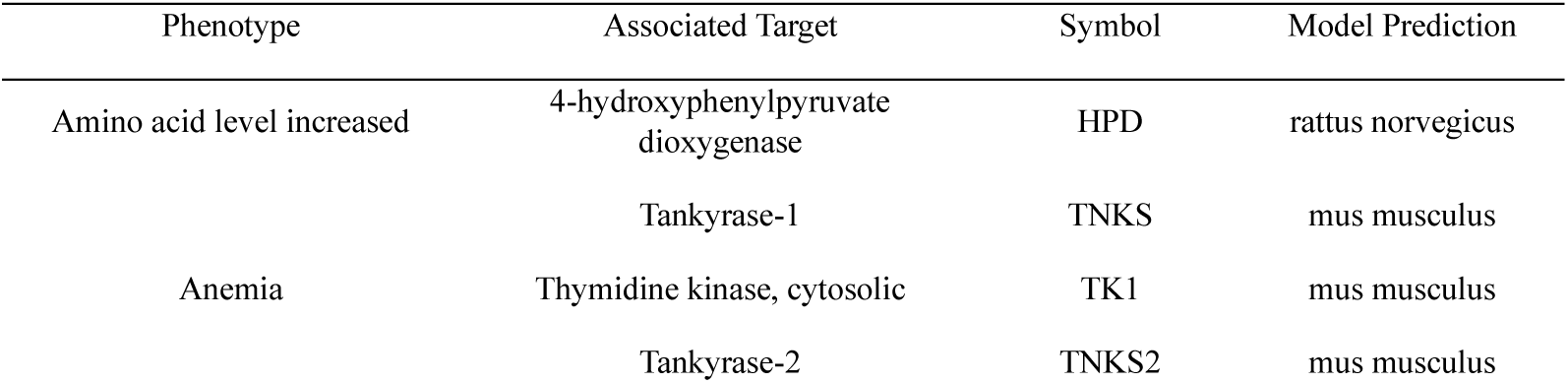
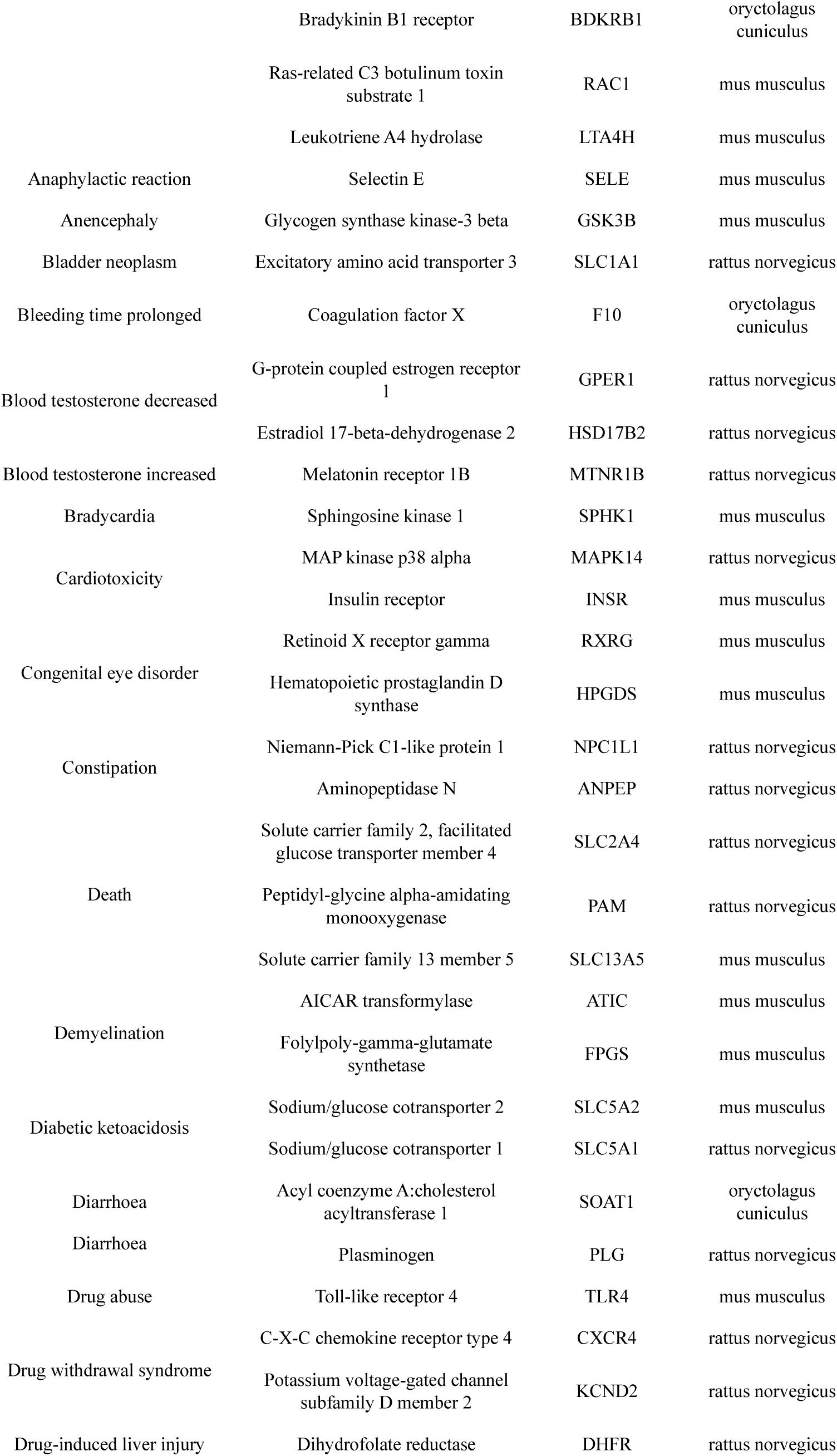
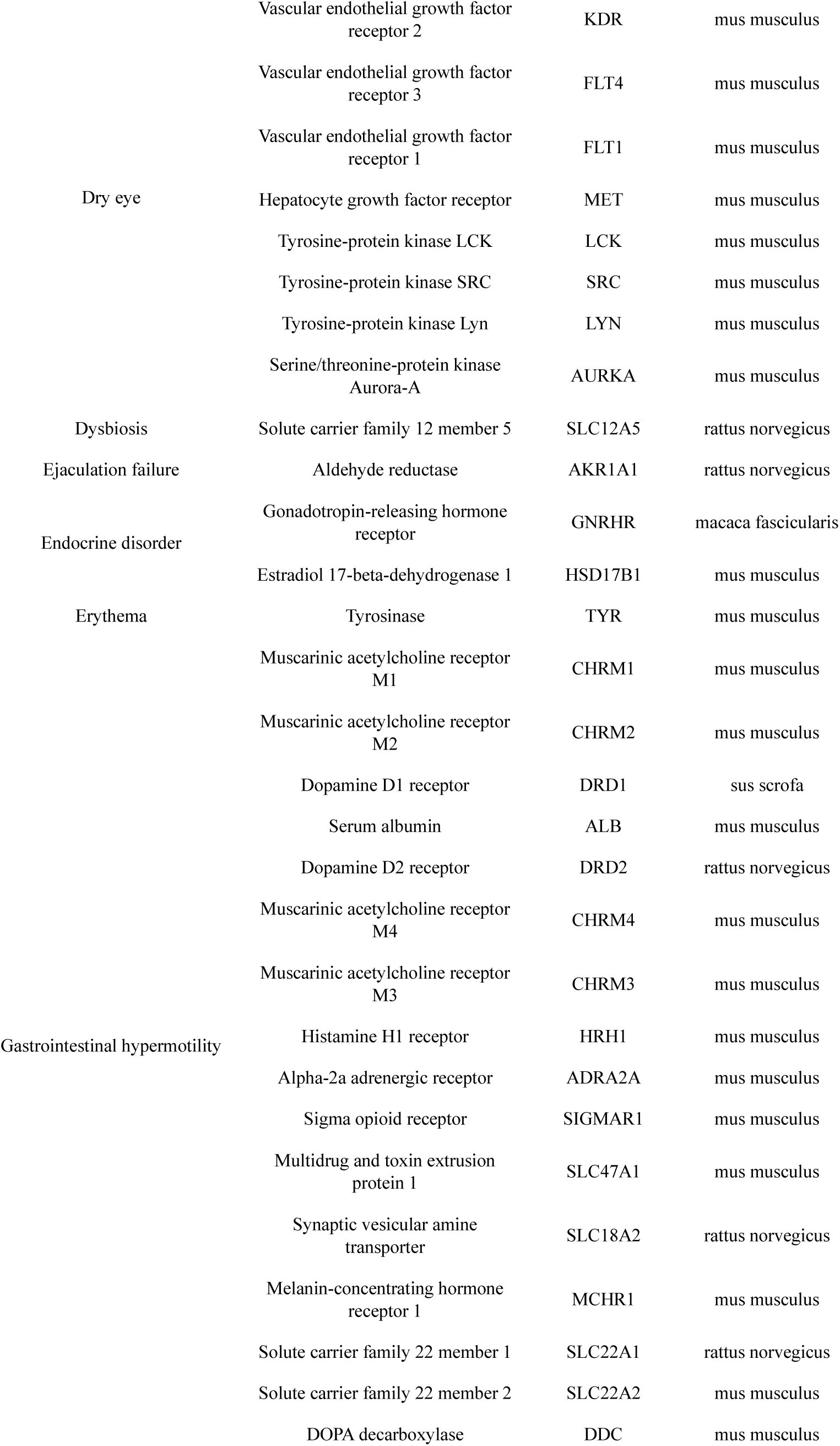
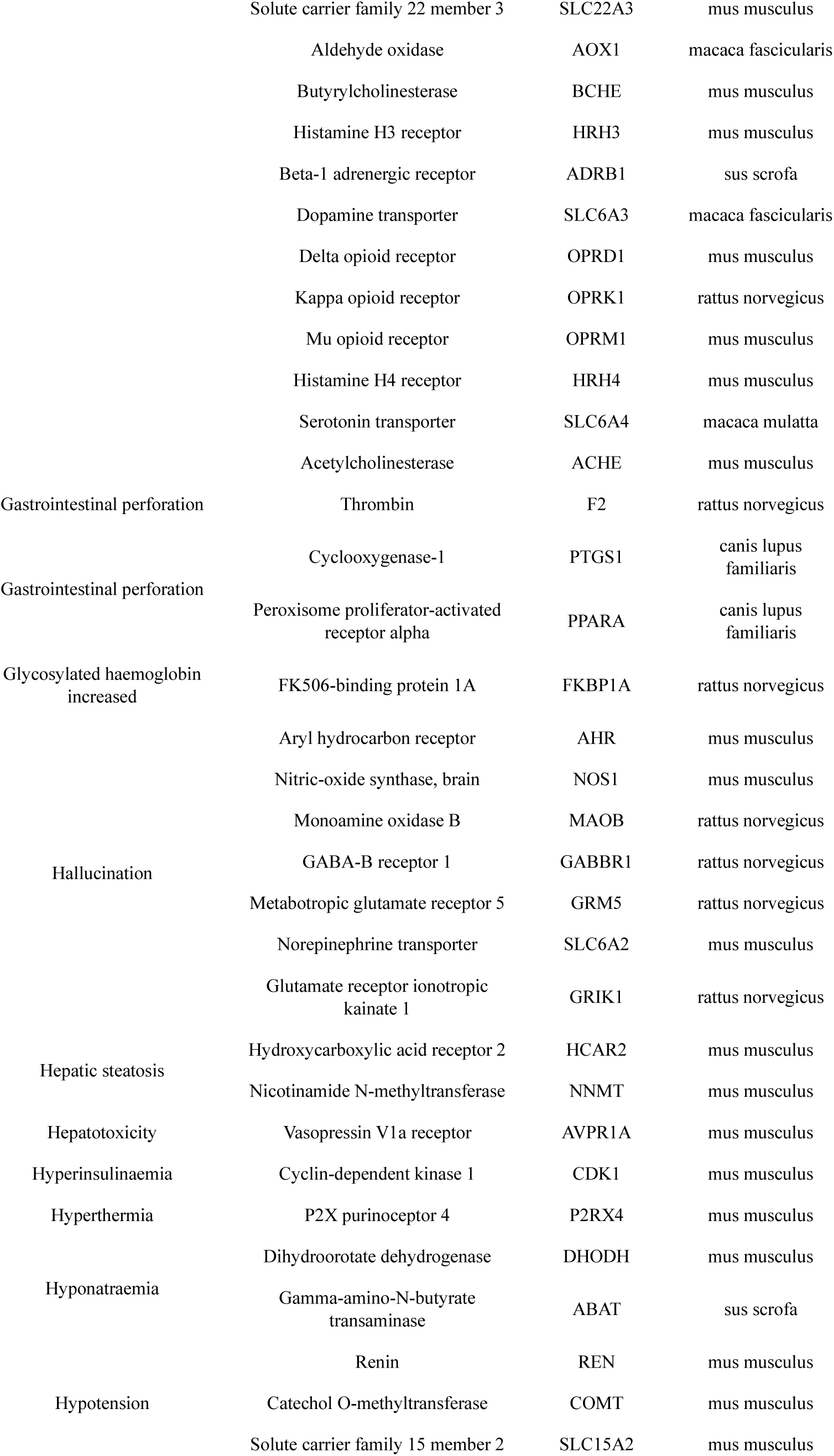
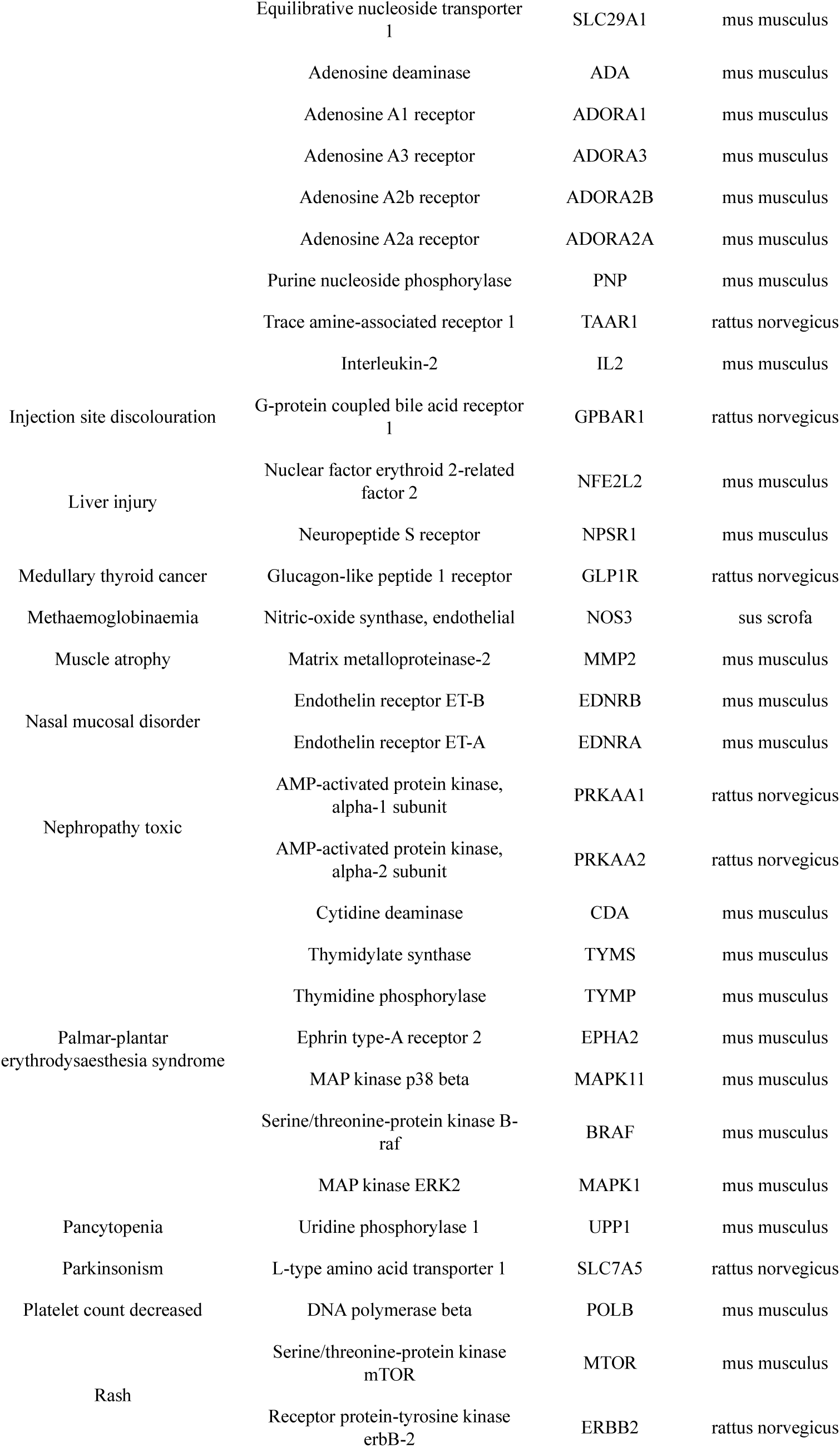
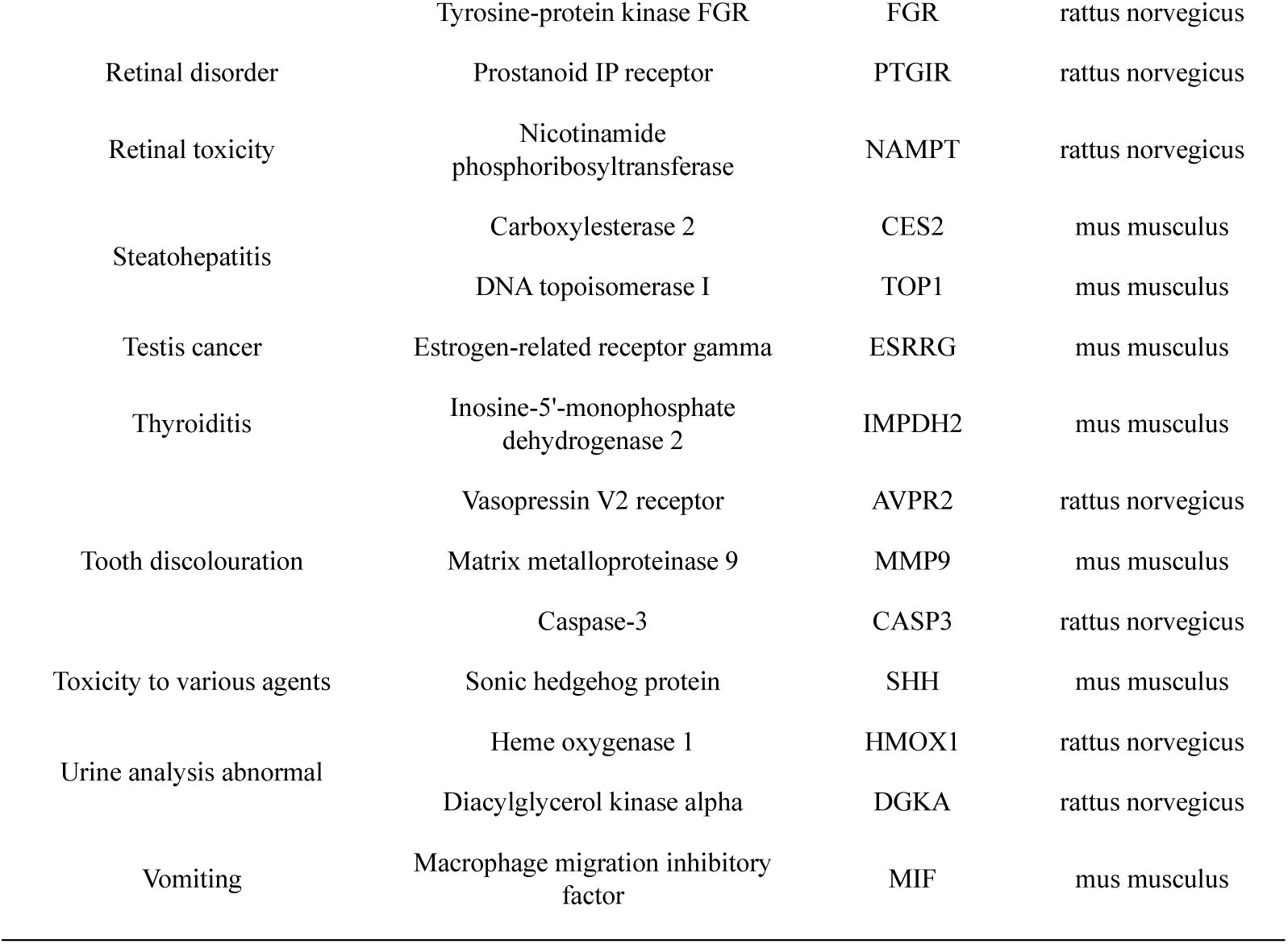
The associated targets for all the phenotypes available in the real-world dataset, as well as the model predictions for those targets are listed below.

